# Screening and testing for a suitable untransfected cell line for SARS-CoV-2 studies

**DOI:** 10.1101/2020.07.09.195040

**Authors:** Claudia Pommerenke, Ulfert Rand, Cord C. Uphoff, Stefan Nagel, Margarete Zaborski, Vivian Hauer, Maren Kaufmann, Corinna Meyer, Sabine A. Denkmann, Peggy Riese, Kathrin Eschke, Yeonsu Kim, Zeljka Macak Safranko, Ivan-Christian Kurolt, Alemka Markotic, Linda Brunotte, Stephan Ludwig, Luka Cicin-Sain, Laura Steenpaß

## Abstract

At present, the novel pandemic coronavirus SARS-CoV-2 is a major global threat to human health and hence demands united research activities at different levels. Finding appropriate cell systems for drug screening and testing molecular interactions of the virus with the host cell is mandatory for drug development and understanding the mechanisms of viral entry and replication. For this, we selected human cell lines represented in the Cancer Cell Line Encyclopedia (CCLE) based on RNA-seq data determined transcript levels of ACE2 and TMPRSS2, two membrane proteins that have been identified to aid SARS-CoV-2 entry into the host cell. mRNA and protein expression of these host factors were verified via RQ-PCR and western blot. We then tested permissiveness of these cell lines towards SARS-CoV-2 infection, cytopathic effect, and viral replication finding limited correlation between receptor expression and infectability. One of the candidate cancer cell lines, the human colon cancer cell line CL-14, tested positive for SARS-CoV-2 infection. Our data argue that SARS-CoV-2 in vitro infection models need careful selection and validation since ACE2/TMPRSS2 receptor expression on its own does not guarantee permissiveness to the virus.

**Author summary:** In the midst of the pandemic outbreak of corona-virus SARS-CoV-2 therapeutics for disease treatment are still to be tested and the virus-host-interactions are to be elucidated. Drug testing and viral studies are commonly conducted with genetically manipulated cells. In order to find a cell model system without genetic modification we screened human cell lines for two proteins known to facilitate entry of SARS-CoV-2. We confirmed and quantified permissiveness of current cell line infection models, but dismissed a number of receptor-positive cell lines that did not support viral replication. Importantly, ACE2/TMPRSS2 co-expression seems to be necessary for viral entry but is not sufficient to predict permissiveness of various cancer cell lines. Moreover, the expression of specific splice variants and the absence of missense mutations of the host factors might hint on successful infection and virus replication of the cell lines.

## Introduction

It is of no debate that the overcoming of the pandemic SARS-CoV-2 pathogen spreading across the continents urgently needs joint efforts in the scientific community. Within a few months in the midst of spring 2020 the pandemia is accompanied with a high case fatality rate of up to 20% for vulnerable risk groups [1] and high excess mortality in Europe monitored by EuroMOMO (https://www.euromomo.eu/graphs-and-maps) [2]. To date no specific antiviral treatment to SARS-CoV-2 has been approved not to mention vaccine treatment. However, promising antiviral drugs such as remdesivir [3], camostat [4], and potential vaccines such as mRNA-1273 [5] are underway. Still, as long as no clinically proven drugs and vaccines are at hand, screening and testing of potential drugs is a major part of the current global research effort as well as studying the molecular mechanisms of viral entry to the host cell. To this, our contribution to the scientific progress is to validate human cell lines of the Leibniz Institute DSMZ for their permissiveness to SARS-CoV-2. Cell lines serve as valuable and valid model systems to study different diseases [6,7] and have been used specifically for SARS-CoV-2 viral entry [4,8] or for antiviral activities against this virus [9]. Particularly the two surface proteins ACE2 and TMPRSS2 have been shown to contribute to viral binding and processing [4,8] and are mainly expressed in bronchial transient secretory cells [10]. Interestingly, high gene expression of ACE2 is shown for other tissues such as myocardial cells, esophagus epithelial cells, and enterocytes, which may hint at COVID-19 patients, infected by SARS-CoV-2, exhibiting non-respiratory symptoms [11–14].

Here, in order to get hold of human cell lines helpful for SARS-CoV-2 studies, we examined human cell lines with comparably high gene expression of ACE2 and TMPRSS2. The rich resource Cancer Cell Line Encyclopedia (CCLE) provided publicly available RNA-Seq data [15], which we screened for human cell lines expressing high levels of these two surface proteins. Selected cell lines were picked for validation on mRNA and protein levels and tested for the ability of SARS-CoV-2 to infect these cells. With these cell lines we devise appropriate, non-overexpression cell model systems to facilitate experimental SARS-CoV-2 work, further the understanding of viral entry and support drug development.

## Results and Discussion

### Selection of cell model systems

Inspired by the publication of Hoffmann et al. pinpointing the host factors ACE2 as SARS-CoV-2 receptor and TMPRSS2 as viral S protein priming serine protease [4], we sought for identifying human cell lines enriched for these two proteins. Fortunately, the Cancer Cell Line Encyclopedia (CCLE) provides large-scale sequencing data including mRNA-seq expression data for over 900 cell lines [15]. Screening for gene expression of ACE2 and TMPRSS2 in this data set enabled us to identify potential susceptible cell cultures prone to SARS-CoV-2 infection. At first, we selected for 301 human cell lines readily in stock at the DSMZ cell lines repository and visualised gene expression of ACE2, which was shown to be the cellular receptor for the viral S protein [16], and TMPRSS2 known to cleave SARS-CoV S protein [17] (Fig 1A). Gene expression for these two proteins differed between and within the various tumour entities. Of these cell lines, the colon carcinoma cell lines CL-14 and CL-40 and the breast carcinoma cell line HCC-1937 belonged to the top 15 ACE2 expressing cell lines along with high expression for TMPRSS2 (Table 1). Since the mucosa of the upper aerodigestive tract shows a high virus replication in vivo [18], we additionally selected the tongue squamous cell carcinoma cell lines CAL-27 and CAL-33, the esophageal squamous cell carcinoma cell line KYSE-510, and the newly established oral squamous cell carcinoma cell lines UPCI-SCC-074 and UPCI-SCC-131. Since the lung carcinoma cell lines Caco-2, Calu-3 and the green monkey kidney cell line Vero E6 are shown to be permissive for SARS-CoV-2 [4, 8], these were used as positive control cell lines.

**Fig 1.**
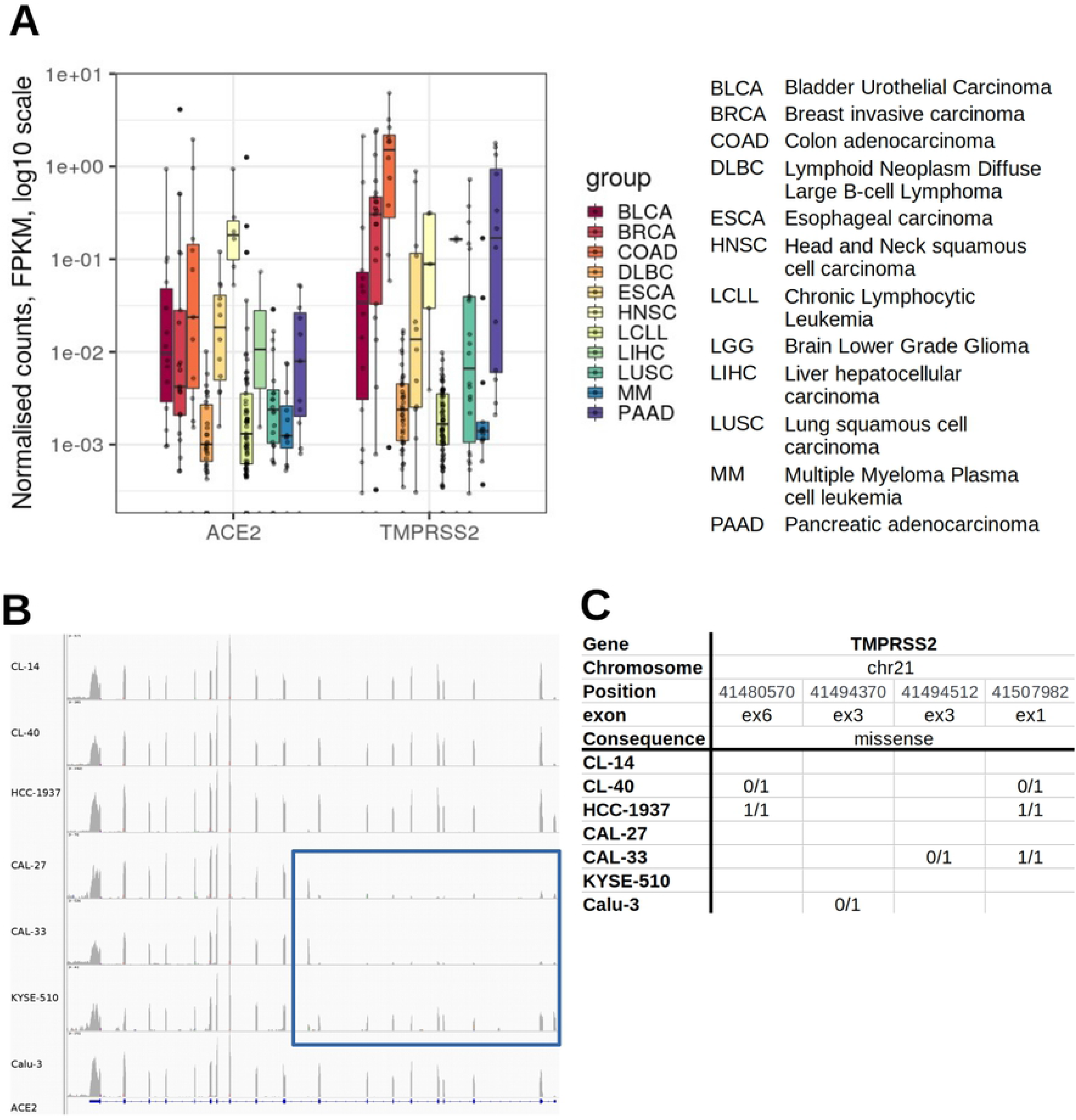
ACE2 and TMPRSS2 gene expression data for 301 DSMZ cell lines in the CCLE RNA-seq data set. (A) Expression of ACE2 and TMPRSS2 for selected disease entities (in logarithmic scale). Gene expression for ACE2 and TMPRSS2 varies between and within the different tumour species. FPKM - fragments per kilobase million: normalised gene expression data. (B) Different exon usage of ACE2 for the selected cell lines. CAL-27, CAL-33 and KYSE-510 show reduced exon expression from ex2 to ex8 (see blue rectangle). ACE2 is reversely oriented. (C) Missense mutations in the coding regions of TMPRSS2 for the selected cell lines. No mutations were found in the coding regions of ACE2. 0/1 denotes heterozygous and 1/1 homozygous mutations.

**Table 1.** Human cell lines exerting high levels of ACE2 and TMPRSS2 based on CCLE RNA-seq data. RNA-seq data were normalised and calculated to FPKM values and the 15 top ACE2 expression cell lines are listed. See Fig 1A legend for group abbreviations. Text coloured in blue list the cell lines tested in this study. FPKM - fragments per kilobase million: normalised gene expression data.

| Cell line | Group | ACE2 | TMPRSS2 |
| --- | --- | --- | --- |
| HCC-1937 | BRCA | 4.126 | 0.468 |
| CL-14 | COAD | 1.956 | 2.646 |
| MOLM-16 | LCLL | 1.255 | 0.001 |
| CL-40 | COAD | 0.958 | 6.204 |
| CAL-33 | HNSC | 0.940 | 0.317 |
| BFTC-905 | BLCA | 0.936 | 0.045 |
| CAL-51 | BRCA | 0.507 | 0.023 |
| HDQ-P1 | BRCA | 0.507 | 0.239 |
| CAL-27 | HNSC | 0.283 | 0.309 |
| M-07e | LCLL | 0.227 | 0.002 |
| SCC-25 | HNSC | 0.198 | 0.000 |
| BHY | HNSC | 0.166 | 0.004 |
| CL-34 | COAD | 0.166 | 3.198 |
| LoVo | COAD | 0.125 | 1.234 |
| KYSE-510 | ESCA | 0.120 | 0.109 |

Of note, gene expression of different splice variants of the ACE2 protein became evident comparing the read numbers of the exons as seen in the CCLE RNA-seq data for CAL-27, CAL-33 and KYSE-510 (Fig 1B). Furthermore, missense mutations were detected in TMPRSS2 for CL-40, HCC-1937, CAL-33 and Calu-3 (Fig 1C). We assume that these variations may have a considerable effect on the virus-host interaction. Recently published data suggest, that cells need interferon stimulation to produce ACE2, thereby changing the cell’s permissiveness to SARS-CoV-2 and hinting on the dynamic levels of this host factor [19] [20].

All selected cell lines exhibited marked ACE2 and TMPRSS2 mRNA expression according to CCLE data or/and RQ-PCR (Table 1 and Fig 2A). Beside qualitative verification for mRNA expression via RNA-Seq data and RQ-PCR (Fig 2A) protein occurrence of ACE2 and TMPRSS2 was confirmed via western blot (Fig 2B).

**Fig 2.**
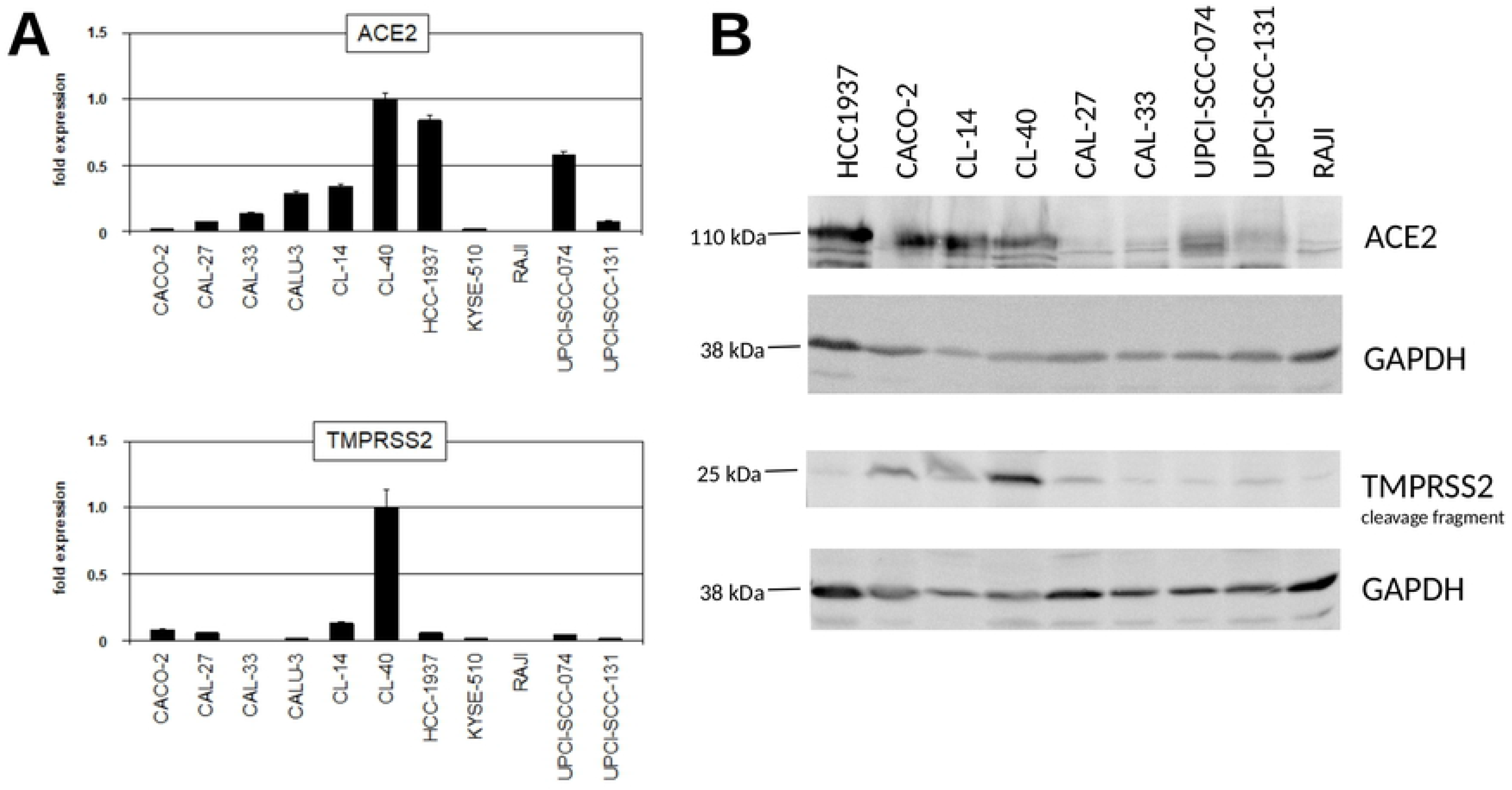
Verified presence of ACE2 and TMPRSS2 in carcinoma cell lines determined by RQ-PCR and western blots. (A) Quantification of ACE2 and TMPRSS2 transcripts in selected cell lines by RQ-PCR. (B) Western blot analysis was performed to assess the expression of proteins associated with the SARS-CoV-2 cell entry. Cell line RAJI served as negative control.

No co-expression of ACE2 and TMPRSS2 was found in the LL-100 data set, another data set which comprises RNA-seq and WES data of 100 human leukemia and lymphoma cell lines [6] (Table S1). This is reflected in the CCLE results, in which cell lines of the hematopoetic lineages exhibited lower gene expression of both genes compared to other cell lines of different origin (see Fig 1A, DLBC, LCLL and MM tumour entities).

The presence of the viral host factors ACE2 and TMPRSS2 in cell lines of various origins is in accordance to previously published data, where ACE2 and TMPRSS2 were detected in cells from multiple tissues including respiratory tract, esophagus and colon [21] [22] [14] [12] and virus entry was evidenced in gastro-intestinal tissues [23].

**Table S1. Human leukemia and lymphoma cell lines express low levels of ACE2 and TMPRSS2 in the LL-100 RNA-seq data.** RNA-seq data were normalised and calculated to FPKM values and ordered to ACE2 expression. FPKM - fragments per kilobase million: normalised gene expression data.

### Testing for SARS-CoV-2 infectivity and permissiveness

Receptor ACE2 and surface proteinase TMPRSS2 have been shown to play a key role for SARS-CoV-2 cellular entry. Whether their co-expression is sufficient to allow entry and whether alternative receptors may exist is still under debate. In order to shed light on infectivity and permissiveness of the appointed cell lines in context of their ACE2/TMPRSS2 expression, we subjected cells to experimental infection with wildtype SARS-CoV-2 virus. Three key parameters of viral infection were quantified: cytopathic effect (CPE), a hallmark of lytic coronavirus infections, viral gene expression, and production of infectious viral progeny.

CPE summarizes a plethora of mechanisms accompanied with lytic viral infection that can be quantified as loss of confluence in cell culture due to cell death [24]. Using time-resolved phase contrast imaging of infected cell lines, we determined time course and extent of SARS-CoV-2-mediated CPE for the cell lines CAL-27, CAL-33, CL-14, CL-40, HCC-1937, KYSE-510, UPCI-SCC-074, and UPCI-SCC-131 (Fig 3A, Fig 3B, data not shown). Cell confluence was reduced after infection for CL-14, Vero E6, and Calu-3 in accordance with previously published data for SARS-CoV [4, 8, 25] and data made available in pre-print [26]. The results show a clear onset of decreasing confluence from two days post infection in Vero E6 cells, the most commonly used cell line to experimentally generate SARS-CoV-2 viral stocks. The same was observed for the lung cancer cell line Calu-3 as well as the colon carcinoma cell line CL-14 but not CL-40, although the ACE2 and TMPRSS2 expression was comparable according to RNA-seq and western blot analysis or indicated even higher levels for CL-40 when determined by RQ-PCR. Patterns of cytopathic disturbance of the cell culture monolayer were specific for each cell line with most drastic effects seen in Calu-3 with a reduction to 80% confluence within 4 days post infection (Fig 3B). Whereas the lacking permissiveness of the selected cell lines CAL-27, CAL-33 and KYSE-510 for SARS-CoV-2 might be explained by a shortened ACE2 transcript (Fig 1B), it is more challenging to elucidate, why the candidate cell line CL-40 tested non-permissive. CL-40 as well as HCC-1937 hold a missense mutation in exon 6 (Fig 1C) causing an amino acid exchange at position 197 (V197M) within the Scavenger receptor cysteine-rich (SRCR) domain responsible for protein-protein interaction and ligand binding (ExPASy) and is described for various cancer types as well as cell lines (COSMIC). Hence, we speculate that this mutation at V197M might disturb interaction with SARS-CoV-2. Antibodies or pharmaceutic agents blocking this region of the receptor protein might be promising candidates in preventing the infection of the cells with SARS-CoV-2.

**Fig 3.**
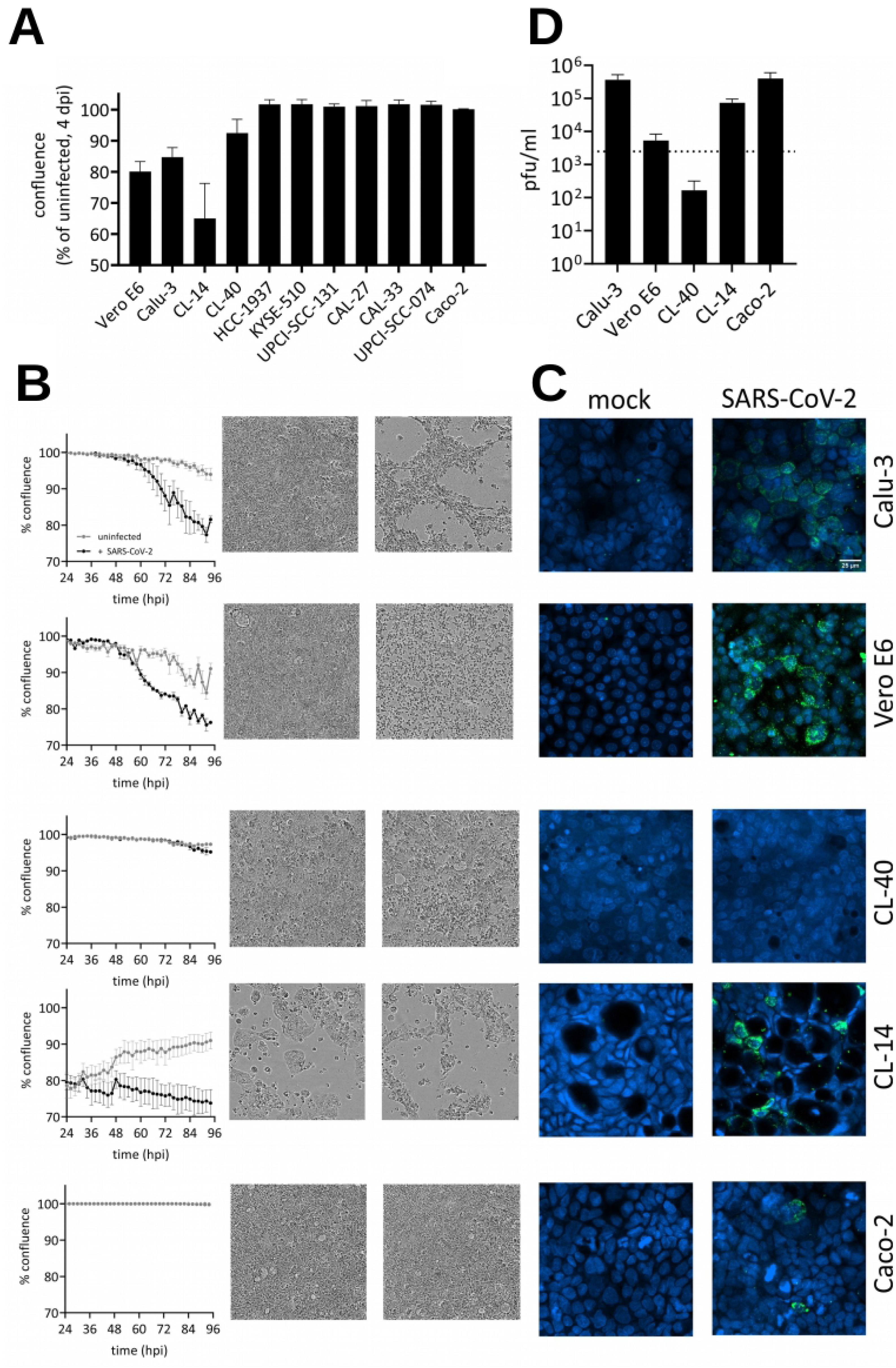
Susceptibility of cell lines toward SARS-CoV-2 infection. (A) Confluence of indicated cell lines at 4 days postinfection with SARS-CoV-2 (strain Zagreb) as percent of the uninfected control. Error bars show SEM of at least three technical replicates. (B) Confluence was determined by phase contrast imaging at 2 h intervals. Representative images show confluence at 1 and 3 dpi. Confluence was determined by the IncuCyte GUI software. Error bars show SEM of three technical replicates. hpi: hours postinfection, dpi: days postinfection. (C) Immunofluorescence staining of viral double-stranded RNA (dsRNA) as indicator for viral replication. Cell lines infected with SARS-CoV-2 or mock-infected were fixed after two days and stained for dsRNA (green) and nuclei (Hoechst 33342; blue). (D) Virus titers in the supernatant of infected cell lines at 5 dpi determined by plaque assay. pfu: plaque-forming units.

Since CPE alone may not correlate directly with the frequency of infection in a cell population, we sought to quantify viral gene expression by immunofluorescence staining viral nucleic acids in form of double-stranded RNA [27] (Fig 3C). Double-stranded (ds)RNA was found in the perinuclear space of infected cells consistent with earlier findings on SARS-CoV [27], indicating active viral transcription and genome replication. Calu-3, Vero E6, and CL-14 cells showed substantial amounts of infected cells with dsRNA in perinuclear regions, but also Caco-2 cells – while showing no cytopathic effect – displayed a limited number of dsRNA-containing cells. CL-40, in contrast, showed no detectable dsRNA-containing cells consistent with the absence of CPE upon infection. Interestingly, Vero E6 showed a more spot-like appearance of dsRNA signals compared to human cells which could reflect a species-specific property of SARS-CoV-2 replication.

Finally, we assessed the permissiveness of these cell lines by assessing the release of mature, infectious virions in a plaque assay (Fig 3D). Supernatants of infected cells were taken 5 days post infection (dpi) and plaques were counted on Vero E6 cells. We found that all tested cell lines, except CL-40, support the production of infectious SARS-CoV-2 progeny to different extents – consistent with the presence of dsRNA in Fig 3C. However, Caco-2 cells produced high amounts of SARS-CoV-2 infectious virions without showing a CPE over the course of five days and also despite of low dsRNA staining.

Cancer cell lines have been indispensable work horses of biological and clinical research, including current global efforts to tackle SARS-CoV-2. Their further careful evaluation as a non-overexpression based SARS-CoV-2 experimental systems is key to future discoveries and their interpretation.

## Conclusion

So far, clinically proven therapeutics specifically addressing SARS-CoV-2 are missing for the treatment of COVID-19 patients. It is getting clear, that the virus attacks different organs and cell types [14,21,22], which explains diverse symptoms of COVID-19 [11–13]. Many virus-host-interaction studies rely on ACE2/TMPRSS2-transfected cell lines such as 293T and HeLa cells or non-human material such as green monkey cells Vero E6 [4,28,29]. By screening RNA-seq data of human cell lines for higher levels of ACE2 and TMPRSS2 gene expression, we selected candidate cell lines, of which the human colon carcinoma cell line CL-14 tested positive for SARS-CoV-2 permissiveness. This non-overexpressing cell line may represent a suitable cell line for SARS-CoV-2 studies. In summary, we find that the expression of ACE2 and TMPRSS2 is necessary but not sufficient for SARS-CoV-2 productive infection in human cell lines. Different ACE2 splicing variants and missense mutations in TMPRSS2 may be a reason for the prevention of infections of receptor expressing cell lines.

## Materials and Methods

### CCLE RNA-Seq data set analysis

RNA-Seq data were retrieved and preprocessed as described [30]. Briefly, alignment files were downloaded from CGHub via genetorrent, sorted (samtools 0.1.19) and converted to fastq files (bedtools v2.21.0). Reads were trimmed via fastq-mcf (ea-utils 1.1.2-686). STAR (2.5.3a) served as read mapper to the human genome Gencode 26, HTSeq (0.11.3) was used as count tool. Raw counts were normalised and transformed to FPKM (fragments per kilobase million) via R/Bioconductor (3.6.9) loading DESeq2 (1.26.0) and visualised via ggplot2 (3.1.1). Different exon usage was discovered by visualising the alignment files via IGV (2.8.0) [31]. For detecting mutations, mapped reads were assigned to read groups (picard, 2.9.2 [www.broadinstitute.github.io/picard]); reads were split, trimmed and reassigned (GenomeAnalysisTK, 3.7-0, SplitNCigarReads [32]); mutations were called by the HaplotypeCaller (GenomeAnalysisTK, 3.7-0,) [32]; and mutation effects were predicted via the Ensembl VEP (GRCh38, v90) [33]. Mutations were filtered ≥10 depth and predicted missense mutations.

### Cultivation of human cell lines

The continuous cell lines were provided for accessioning to the biological resource center DSMZ by the original or secondary investigators [34]. Cell lines were grown at 37’C in a humidified atmosphere of air containing 5% CO2. The basic growth media (Life Technologies, Darmstadt, Germany) were supplemented with 10–20% fetal bovine serum (Sigma Aldrich, Taufkirchen, Germany). No antibiotics were added to the cultures. All cell lines were free of mycoplasmas and other bacterial, yeast and fungi contaminations as tested by PCR and microbiological growth assays [35]. The authenticity of the cell lines was determined by DNA typing [36]. The cells were first cultured in cell culture flasks to confluence, detached with trypsin/EDTA, washed with PBS and pelleted for RNA and protein preparation. For virus infection experiments the cells were seeded in 96-well plates at different densities starting with 0.25 – 1.5 x 105 cells per well as highest cell number depending on the cell line. Optimal cell densities were determined by observing the growth progression with an IncuCyte instrument (Essen Bioscience, Hertfordshire, UK). Cells were then seeded at semi-confluence or complete confluence as highest cell densities and three further dilutions each with half of the previous cell densities. The cells were grown for one day before the virus infection experiments were started.

### RQ-PCR analysis

Real-time quantitative (RQ) polymerase chain reaction (PCR) analysis was performed to quantify specific transcripts in the cells. Total RNA was extracted from cell line samples using TRIzol reagent (Fisher Scientific, Schwerte, Germany). cDNA was generated from 1 μg RNA by random priming using the Biozym cDNA Synthesis Kit (Biozym, Hessisch Oldendorf, Germany). RQ-PCR analysis was performed with the 7500 Real-time System, using commercial buffer and primer sets for the amplification of ACE2 (*Hs*01085333_*m*_1) and TMPRSS2 (*Hs*01122322_*m*_1) transcripts (Thermo Fisher Scientific, Darmstadt, Germany). We analysed the transcript of TATA box binding protein (TBP) for normalization of expression levels. Quantitative analyses were performed in triplicate.

### Western blots

Antibodies against ACE2 (ab15348), TMPRSS2 (ab109131), and GAPDH (ab8245) were obtained from Abcam (Cambridge, UK). Western blot samples were prepared as described previously [8]. Proteins on nitrocellulose membranes were visualized with the biotin/streptavidin-horseradish peroxidase system (GE Healthcare; Little Chalfont, UK) in combination with the “Renaissance Western Blot Chemoluminescence Reagent” (Perkin Elmer; Waltham, MA, USA). Documentation was performed using the digital system ChemoStar Imager (INTAS, Göttingen, Germany).

### Viral infection of cell lines

Infection with SARS-CoV-2 was performed using two different virus strains: Zagreb (SARS-CoV2/ZG/297-20, University Hospital for Infectious Diseases, Zagreb, Croatia), Ischgl (hCoV19/Germany/FI1103201/2020, GISAID accession EPIISL_463008 was taken from the strain repository of the Institute of Virology, Munster). If not indicated otherwise, cells were seeded onto 96-well cell culture plates and infected 24-48 hours later with 250 pfu/well using a 10 *μ*l inoculum and 100 *μ*l total well volume. In order to monitor disturbances of the cell monolayer, images were taken automatically every 2 hours following infection using a Sartorius IncuCyte S3 (10x magnification, 4 images per well) for a total of 7 days. Changes in confluence were analysed using the IncuCyte GUI-inbuilt analysis tools (version 2019B, Rev1).

### Virus plaque assay

Quantification of SARS-CoV-2 infectious units was done via a virus plaque assay. Supernatants of infected cells were taken 5 dpi, serially diluted, and used to infect confluent Vero E6 cells on 96-well cell culture plates for one hour. Then, the inoculum was removed and cells were overlaid with cell culture medium (MEM, 10% FCS, 2 mM glutamine) containing 1.5% methyl-cellulose. After 3 days, virus plaques were counted from phase contrast microscopic images.

### Immunofluorescence staining

SARS-CoV-2 infected cells were fixed with 6% paraformaldehyde in PBS for one hour at room temperature, followed by washing three times with PBS. Cell permeabilisation was done with PBS containing 0.15% Triton X-100 for 10 min at RT. Primary antibody used to stain dsRNA was mouse anti-dsRNA, clone rJ2 (Merck Millipore, Cat. no. MABE1134); secondary antibody was goat anti-rabbit-Cy5 (Life Technologies, Cat. no. A10523).

## Acknowledgments

We thank Hilmar Quentmeier for initial coordination of activities for this project. This project was supported by the grant 14-76103-184 from the Ministry of Science and Culture of Lower Saxony to Luka Cicin-Sain.

## Author Contributions

Methodology: CP, CCU, UR; Investigation: UR, MZ, SN, VH, MK, SAD, CP, PR, KE, YK; Formal Analysis: CP, UR, SN, CCU; Resources: ZMS, ICK, LB, ZML, ICK, AM, LB, SL; Writing - Original Draft: CP, UR, CU; All authors have read and approved the final manuscript.

